# Label-free characterization of visual cortical areas in awake mice via three-photon microscopy reveals correlations between functional maps and structural substrates

**DOI:** 10.1101/790436

**Authors:** Murat Yildirim, Ming Hu, Peter T. C. So, Mriganka Sur

## Abstract

Our understanding of the relationship between the structure and function of the intact brain is mainly shaped by magnetic resonance imaging. However, high resolution and deep-tissue imaging modalities are required to capture the subcellular relationship between structure and function, particularly in awake conditions. Here, we utilized a custom-made three-photon microscope to perform label-free third-harmonic generation (THG) microscopy as well as laser ablation to calculate effective attenuation lengths (EAL) of primary visual cortex and five adjacent visual cortical areas in awake mice. We identified each visual area precisely by retinotopic mapping via one-photon imaging of the calcium indicator GCaMP6s. EALs measured by depth-resolved THG microscopy in the cortex and white matter showed correspondence with the functional retinotopic sign map of each cortical area. To examine the basis for this correspondence, we used THG microscopy to examine several structural features of each visual area, including their cytoarchitecture, myeloarchitecture and blood vessel architecture. The cytoarchitecture of each area allowed us to estimate EAL values, which were comparable to experimental EAL values. The orientation of blood vessels and myelin fibers in the six areas were correlated with their EAL values. Ablation experiments, which provide ground truth measurements, generated 17 ± 3 % longer EALs compared to those obtained with THG imaging but were consistent with the latter. These results demonstrate a strong correlation between structural substrates of visual cortical areas, represented by EALs, and their functional visual field representation maps.

## Introduction

An important question in neuroscience is to understand the relationship between the structure and function of brain regions because of the close relationship between the two^1^. The structure and function of brain regions can be defined with various metrics. Cytoarchitectonics^2^, and myeloarchitectonics^3,4^ are commonly used metrics for defining structural features of the brain. Resting state or spontaneous activity^5,6^, neuronal activity evoked by a sensory stimulus^7^ or during behavior^8^, and retinotopic mapping^9^ are the most established metrics for defining the function of brain regions. Previous studies have shown the relationship between cytoarchitecture and function^10,11^, as well as between myeloarchitecture and function of brain regions^12–14^ among humans and nonhuman primates. These relationships can be measured by structural and functional magnetic resonance imaging (MRI)^15^. Since the spatial resolution of MRI is on the order of 100 μm at best^16^, it does not provide adequate measurement resolution for many brain regions in small animals such as mice. Therefore, high-resolution in vivo imaging technologies are required to relate the structure and function of the brain of small animal models.

Multiphoton microscopy has revolutionized neuroscience by providing subcellular resolution as well as capabilities for deep-brain imaging^17–19^ in mice. However, there are limitations to both two- and three-photon microscopy with respect to maximum imaging depth, depending on signal-to-noise ratio, as well as optical properties such as scattering and absorption lengths of specific brain regions^19,20^. Several of these features are related to structural components of the cortex, such as cytoarchitecture and myeloarchitecture^21,22^. The effective attenuation length (EAL), defined as the combination of scattering and absorption lengths, provides a correlate of these structural components and is crucial for determining maximum imaging depth of any brain region for performing reliable imaging and manipulation of cells^23–25^. The most common technique to determine the EAL is by labeling blood vessels via the injection of high absorption cross-section dyes retro-orbitally for two- and three-photon microscopy in anesthetized mice^26^. However, this method assumes uniform labeling of blood vessels, as well as similar optical properties of the anesthetized and awake animal brain, and is based on both excitation and emission photons. Limitations of these assumptions will affect the precision of this technique for calculating the EAL. Due to these limitations, as well as the limited duration of the dye in the blood vessels after retro-orbital injection, alternative and more precise methods are required to characterize the EALs for multiple brain regions in the same animal.

We recently showed that the EAL can be estimated either with third-harmonic generation (THG) imaging of blood vessels and myelin fibers at 1300 nm excitation wavelength in the cortex and in the white matter, respectively, or focally ablating tissue at multiple depths in the primary visual cortex (V1) of awake mice^19^. THG imaging is a label-free three-photon microscopy technique that is sensitive to index of refraction and third order susceptibility changes^27,28^. Previous studies have shown that THG could be used to image blood vessels^19,29^, neurons and their processes^30,31^, and myelin^22^ in the vertebrate central nervous system. Thus, THG imaging of blood vessels and myelin fibers in awake mouse brain can provide in vivo structural information. Together with accurate functional identification of brain regions, such as by using retinotopic maps, THG imaging can then be used to relate their structure to their function.

Cortical areas in the mouse brain can be broadly localized via their stereotaxic coordinates^32,33^; this method is effective for targeting large sections of cortex such as primary visual cortex (V1), motor cortex (M1), or somatosensory cortex (S1). However, this method is less accurate when used to target small functional regions, such as higher visual areas in mice (for example, the anterolateral (AL), lateromedial (LM), posteromedial (PM), anteromedial (AM), rostrolateral (RL) regions of visual cortex) as identified by widefield imaging of intrinsic optical signals^34,35^ or imaging of calcium indicator activity such as GCaMP6s^36^. In these imaging methods, thin line stimuli (often with flickering black-and-white checkerboard segmentation) are drifted in the azimuth and elevation directions to identify retinotopic representations and importantly the boundaries of these visual areas via the generation of a visual retinotopic sign map^35,37^. Primary visual cortex in mice can in some instances be colocalized with cytoarchitecture, chemoarchitecture, and myeloarchitecture features^36,38,39^. However, it has been difficult to find unique structural markers for these areas, and their identification remains solely based on retinotopy and even sign map alone. Importantly, there have been no detailed *in vivo* studies relating structural and functional properties of these visual areas, and in awake mice.

Here, we identified six visual areas (V1, PM, AM, RL, AL and LM) through retinotopic mapping via one-photon imaging of GCaMP6s transgenic awake mice. Next, we performed depth-resolved THG imaging in each area to determine their corresponding EALs in the cortex and in the white matter. We find that these EALs are somewhat unique to each area and are correlated with the retinotopic sign map of visual areas. We next analyzed the cytoarchitecture, in vivo myeloarchitecture, and blood-vessel architecture in each area. The cytoarchitecture of each area allowed us to estimate their EAL values, which we found are comparable with experimental EAL values. Moreover, the orientation of blood vessels and myelin fibers in each area showed similar trends as their EAL values. Finally, we found that ablation experiments, which enable ground truth measurements, provide 17 ± 3% higher EALs compared to those obtained with THG imaging, but are consistent with THG measurements. Thus, THG imaging of blood vessels and myelin fibers provide valid EAL values in the brain. Overall, these results suggest that there is a strong correlation between structure and function of mouse visual areas when we compare their EALs with their retinotopic sign map using sub-cellular resolution label-free THG imaging and functional mapping in awake mice.

## Results

### Calculation of effective attenuation lengths via two label-free methods in six visual areas

We first determined the retinotopic mapping of cortical areas for each animal with the use of one-photon microscopy in mice expressing GCaMP6s (Fig. 1A, see Methods and Materials for details). By drifting a bar containing a checkerboard visual stimulus from −π to π degrees in both azimuthal and altitude directions, we could reliably distinguish six visual areas – V1, PM, AM, RL, AL, and LM (Fig. 1B-D). Taking the blood vessel architecture and retinotopic map as a reference, we utilized a custom-made three-photon microscope^19^ (Fig. 1E-F) to perform depth-resolved THG imaging at 1300 nm in each cortical region, including the cortex and white matter spanning >1 mm thickness (Fig. 1G). Then, we characterized the attenuation of THG signal with respect to imaging depth for each region to calculate their EAL for the cortex and the white matter (Fig. 1H). Larger slopes represent lower EAL (such as in area RL), whereas smaller slopes represent higher EAL (such as in area PM). We calculated three EAL values for each area (cortex-only, white matter-only, and the combination of the cortex and the white matter) in four awake mice. All of these EAL values showed similar trends across all animals (Fig. 1I-K). PM and AM had the highest EAL values in the cortex compared to other visual areas, whereas LM and RL had the smallest EAL values in the cortex (Fig. 1I; see Fig. 3 for statistical comparisons). When we considered EAL values in the white matter, LM, RL, and PM had lower EAL values than those in V1, AL, and AM (Fig. 1J). Finally, when we combined EAL values in the cortex and the white matter as a single EAL value (EAL_comb_), LM and RL had still lower EAL values compared to those in other areas (Fig. 1K). Thus, these results indicate systematic changes in EAL values between cortical areas and representations.

**Figure 1.**
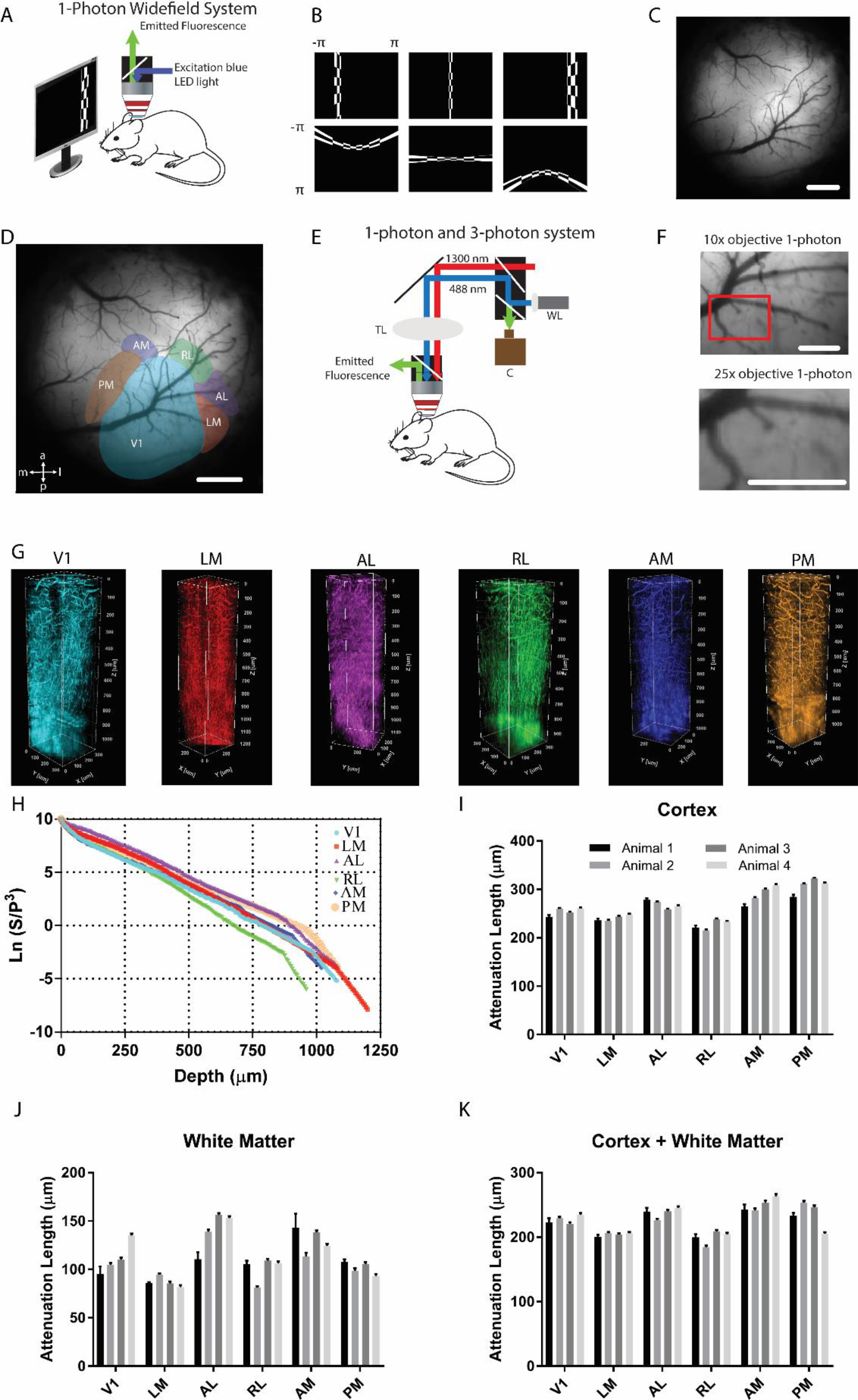
(A) One-photon imaging system to obtain retinotopic maps of visual areas. Blue light was used to excite GCaMP6s in the brain of awake head-fixed mice. Emitted light (green) was collected by the camera. (B) Vertical and horizontal bars (containing flickering black and white checkerboards, and corrected for screen distance and curvature) were drifted across the screen and used to map visual areas of the cortex. (C) Blood vessel architecture of an example mouse with cranial window in the right hemisphere. (D) Representation of six cortical visual areas overlaid on the blood vessel architecture. (E) Combined one- and three-photon system to perform depth-resolved THG imaging. (F) Low magnification (10x) and high magnification (25x) images of V1 obtained with one-photon imaging system coupled with three-photon imaging system. (G) Three-dimensional rendering of a sequence of >200 lateral THG images acquired with 5-μm increment. THG signal is generated in the blood vessels in the cortex and is generated in myelin fibers in the white matter. Each color represents a different visual area. (H) Semi-logarithmic plot for ratio of PMT signal (S) and cube of laser power (P) with respect to imaging depth for third harmonic generation (THG) imaging. Slope of this curve provides EAL values for the cortex and for the white matter. As the slope increases (decreases), EAL value decreases (increases). EAL in the cortex (I), in the white matter (J) and combination of the cortex and the white matter (K) shows similar trends across four awake mice. Scale bars present 1 mm in (C-D), and 0.5 mm in (F).

To evaluate whether THG imaging of blood vessels and myelin fibers at 1300 nm can be used to effectively estimate EAL values, we performed ablation to four depths in the same cortical column in each cortical area. For example, we compared our THG imaging results with ablation results in V1 (Fig. 2). First, we calculated EAL values in the cortex and in the white matter by characterizing the attenuation of THG signal at 1300 nm (Fig. 2A). The slopes of these curves resulted in 259.7 ± 14.6 μm EAL for the cortex and 104.5 ± 4.4 μm EAL for the white matter. To apply the ablation method in awake mouse cortex, four different depths ranging from 150 to 600 μm in 150 μm increments were ablated. A characteristic plot showing percent ablation versus laser energy at brain surface (Fig. 2B) for 150 μm ablation depth showed that 30.0 nJ pulse energy on the surface was required to obtain 50% ablation, where the targeted ablation diameter was 25 μm. To calculate the percent of ablation damage, GCaMP6s and THG images were obtained before and after ablation with varying pulse energies from 10 to 70 nJ. The slope of the threshold energy (*E*_th_) versus ablation depth (Fig. 2C) corresponded to 299.6 ± 17.4 μm extinction length, while the y-axis intercept provided a threshold fluence of 1.44 ± 0.13 J/cm^2^. We also performed ablation experiments in other visual areas (see Supplementary Figures 1–5) and concluded that EAL values obtained with ablation experiments are 17 ± 3 % higher than those obtained with THG imaging (Fig. 2D). Importantly, the relative EAL values were highly consistent between the two measurements. This comparison also showed the clustering of EAL values of medial (PM, AM) areas and lateral areas (RL, LM, and AL) (Fig. 2D). Medial regions had higher EAL values (> 300 μm for THG, >350 μm for ablation) than those in lateral areas (<275 μm for THG, and <300 μm for ablation).

**Figure 2.**
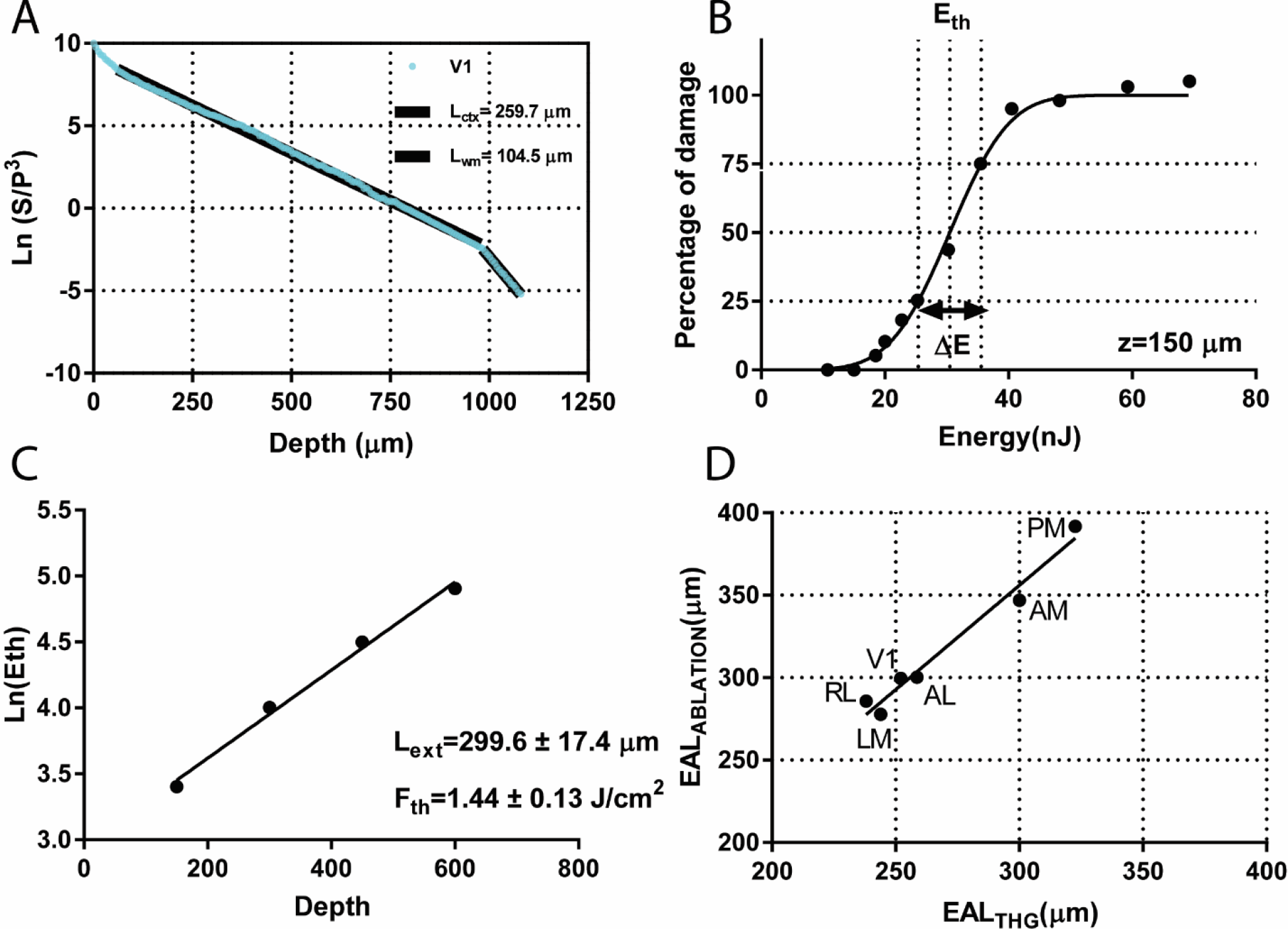
Characterization of EAL values via THG imaging and ablation in V1. (A) THG imaging in V1 results in 259.7 ± 14.6 μm effective attenuation length for the cortex and 104.5 ± 4.4 μm effective attenuation length for the white matter. (B) Determining extinction length via tissue ablation. Percent of damage ranges from 0 to 100% with respect to laser energy on the tissue surface. Threshold energy (*E*_th_) is the energy which results in 50% damage. For ablation at 150 μm depth, *E*_th_ is 30.0 nJ. (C) Semi-logarithmic plot of threshold energies for 4 different depths results in attenuation length of 299.6 ± 17.4 μm and threshold fluence of 1.44 ± 0.13 J/cm^2^. (D) Comparison of EAL values obtained with THG imaging and ablation. Lateral cortical regions such as LM, AL, and RL have lower EAL values compared to those in medial regions such as AM and PM.

### EAL values of visual areas exhibit similar trends as retinotopic sign maps

The boundaries of visual areas were determined by generating their visual field sign map where each visual area has a modulo sign value of −1 or 1. In this sign map, each region has an opposite sign compared to the sign of the regions adjacent to it and with which it shares a significant border – flipped signs reflect mirror-image representations in adjacent areas with a common border^7,36,37^. Thus, the sign of V1 is reversed in its adjacent regions with a significant border (LM, RL, PM). AL has a flipped sign with respect to LM and RL, whereas AM has a flipped sign with respect to PM. These sign shifts can be arbitrarily represented in order as: V1 (+), LM (−), AL (+), RL (−), AM (+), and PM (−). We examined whether our EAL values were unique to each region and significantly different from regions next to each other (Fig. 3). First, we found that V1 has a significantly different EAL value than any other region (Fig. 3A). Many of the other regions also had different EAL values compared to other regions. For example, LM has a significantly different EAL than other regions except RL (Fig. 3A, right). Thus, the EAL is potentially a unique marker of each visual cortical area. Importantly, we observed significant differences in EAL values between regions adjacent to each other in the order mentioned above (except AM and PM) (Fig. 3A, right). Furthermore, the EAL values showed the same trend as the sign maps, so that areas with positive sign had higher EALs than areas with negative sign (Fig. 3A, left). The trend was particularly apparent when we compared EAL values in regions next to each other. The only exception occurred between AM and PM where AM had a lower EAL value than PM (Fig. 3A, left), which may relate to the quite medial location of these areas in the cortex.

**Figure 3.**
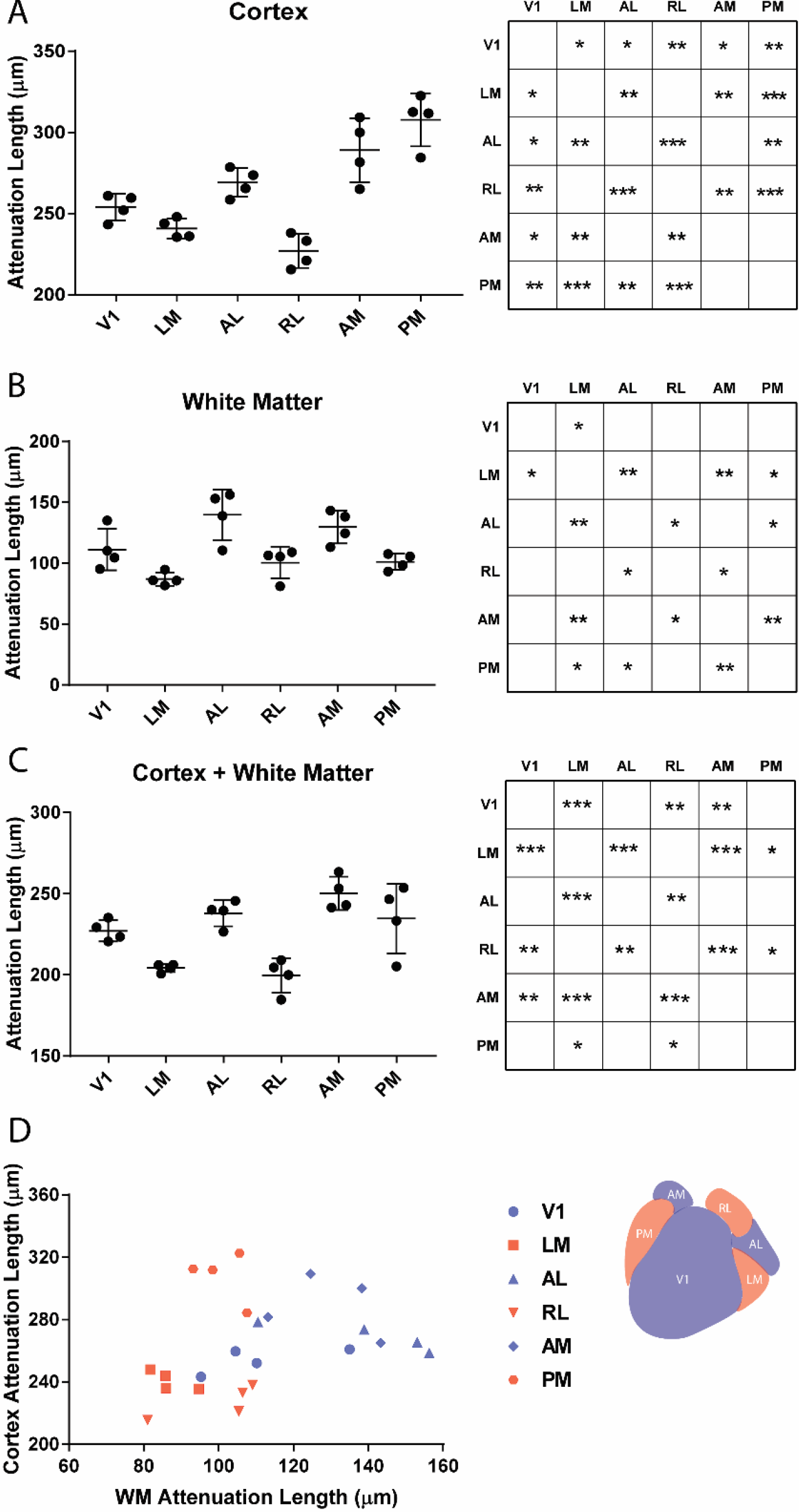
Comparison of EAL values via THG imaging across visual areas in four animals. (A) Comparison of EAL values in the cortex of visual areas (left); p values of one-way ANOVA test with multiple comparisons for EAL values of cortex across all visual areas (right). * represents p<0.05, ** represents p<0.01, and *** represents p<0.001. (B) Comparison of EAL values in the white matter of visual areas (left); p values of one-way ANOVA test with multiple comparisons for EAL values of the white matter across all visual areas (right). (C) Comparison of EAL values of combined cortex and white matter of visual areas (left); p values of one-way ANOVA test with multiple comparisons for EAL values of the combination of cortex and white matter across all visual areas (right). (D) Comparison of EAL values via THG imaging in the cortex and in the white matter across visual areas in four animals. Scatter plot of EAL values in the cortex and in the white matter of six visual areas provides clustering of these regions. Inset: Retinotopic map of six visual areas where the sign of each region is represented by either blue or red. Blue represents three regions (V1, AM, AL) with + sign; red represents three regions (PM, RL, LM) with − sign.

Next, we compared EAL values in the white matter and found that there was a significant difference in EAL values between regions next to each other (Fig. 3B, right). Importantly, there was a continuous decrease and increase in EAL values from V1 to PM. In other words, V1, AL, and AM had significantly higher EAL values than LM, RL, and PM, respectively (Fig. 3B, left). These changes thus followed a similar trend as the functional retinotopic sign map in all six cortical areas. However, there was an opposite relationship between EAL values of AM and PM in the cortex and in the white matter.

We compared the combined EAL values of the cortex (see Methods and Materials) and the white matter. There was a significant difference in EAL values between regions next to each other except between AM and PM (Fig. 3C, right). More importantly, there was a continuous shift in the gradient of these EAL values from V1 to PM (Fig. 3C, left). The average EAL of the cortex-only, white matter-only, and combination of the cortex and white matter for the six visual areas are tabulated in Table 1.

**Table 1.** Mean effective attenuation lengths (EAL) for cortex-only, white-matter-only, and combination of cortex and white matter across four awake mice

| **Length/Area** | **V1** | **LM** | **AL** | **RL** | **AM** | **PM** | **S1** |
| --- | --- | --- | --- | --- | --- | --- | --- |
| **EAL<sub>ctx</sub> (μm)** | 254 | 240.8 | 269.1 | 227.0 | 289 | 307.8 | 312.4 |
| **EAL<sub>wm</sub> (μm)** | 111.2 | 87.1 | 139.7 | 100.5 | 129.9 | 101.2 | 105.4 |
| **EAL<sub>comb</sub> (μm)** | 227.1 | 204.3 | 237.9 | 199.5 | 250.2 | 234.7 | 253.8 |

Finally, we compared attenuation lengths of each region in the cortex and in the white matter for each animal (Fig. 3D). This comparison demonstrated clustering of brain regions with the same sign in the retinotopic map: LM, RL, and PM formed a cluster with low white matter EAL values whereas AL and AM formed a cluster with high white matter EAL values. V1 was located between these two clusters.

### Cytoarchitecture, myeloarchitecture, and blood vessel architecture of visual areas can estimate their EAL values

We first characterized the cell size and cell density of these visual areas in order to describe their cytoarchitecture. For this purpose, we sacrificed each animal after three-photon measurements, sliced their brains, and registered each coronal sections to the Allen Brain Atlas using a custom algorithm^40^. With this algorithm, we could compensate any errors due to tilt of the coronal sections in low-magnification images. Then, we performed high-magnification imaging in individual visual areas (Fig. 4A) and quantified the average cell size and cell density (see Methods and Materials) (Fig. 4B). We estimated the scattering length of each region by using their average cell size and density values via a numerical model^21^ (see Methods and Materials). To estimate the EAL values for each region, we combined the scattering lengths with the absorption length of water at 1300 nm^41^. Comparing the estimated EAL values with the experimental values for the same mouse (Fig. 4C) showed that the estimate of EAL values from the cytoarchitecture of these visual areas agreed well with experimental EAL values obtained with THG imaging in the cortex.

**Figure 4.**
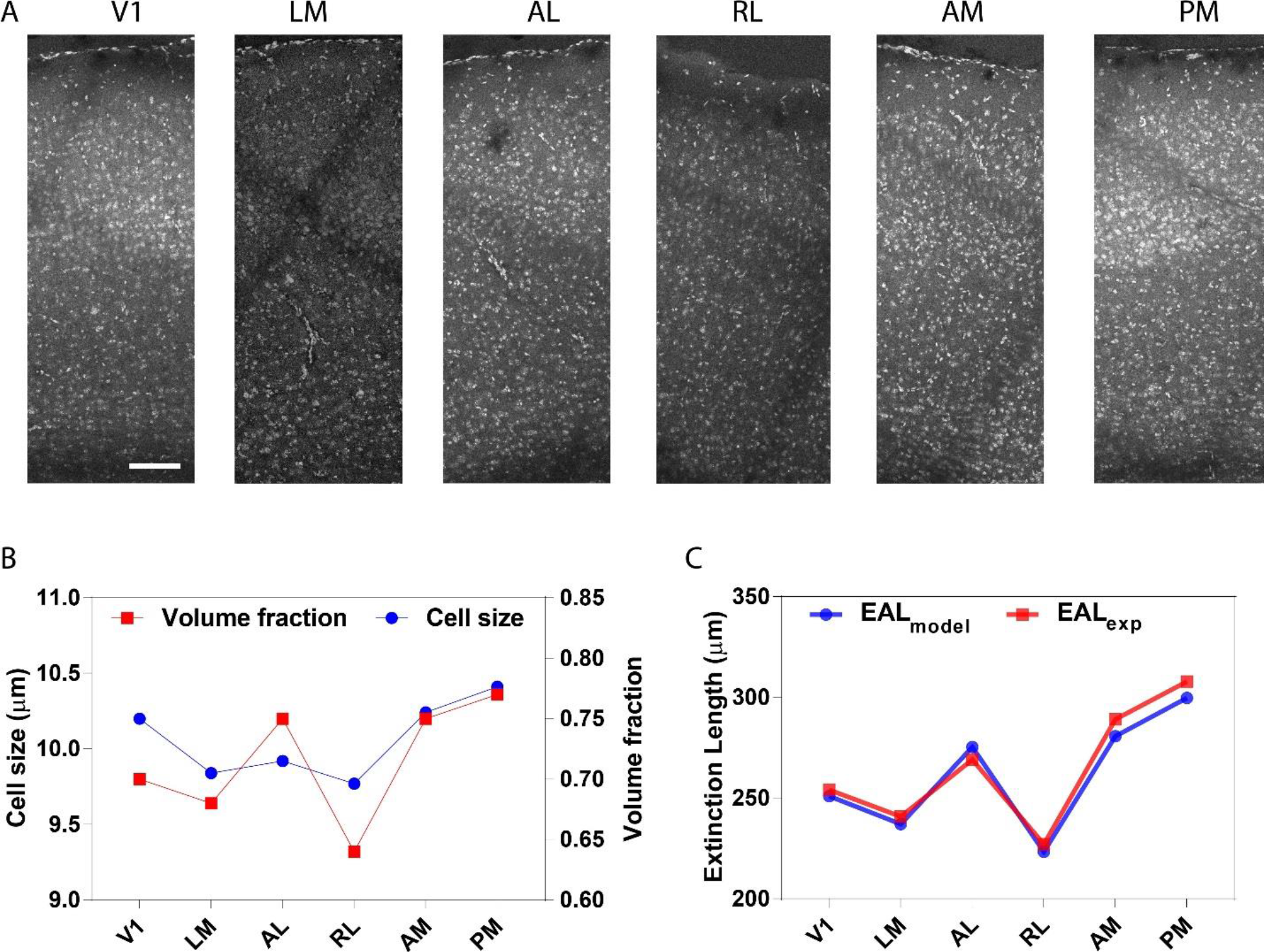
Quantification of cell size and volume fraction, and estimate of effective attenuation length (EAL). (A) High magnification images of cortical column cytoarchitecture of the six cortical areas. (B) Quantification of average cell size and volume fraction for all six visual areas. (C) Comparison of experimental and theoretical EAL values.

We also investigated the *in vivo* myeloarchitecture and blood vessel architecture of these visual areas to see whether they were correlated with EAL values and the retinotopic sign map. We performed depth-resolved THG imaging with finer step size (1 μm) in the axial direction to calculate the orientation of blood vessels and myelin fibers with respect to the same horizontal direction in the cortex and in the white matter (Fig 5A), respectively. First, we analyzed the orientation of the blood vessels in the cortex and demonstrated the distribution of their orientation in polar plots for each region (Fig. 5B). To determine whether the distribution of blood vessel orientation in each visual area was significantly different from others, we applied a circular multi-sample one-factor ANOVA statistical test (Fig. 5C) (see Methods and Materials). This analysis revealed that blood vessel orientation distributions in AM and PM were significantly different than orientation distributions in V1, LM, RL, and AL. Furthermore, we calculated the circular mean of these orientation distributions and found that circular mean values of AM and PM regions were higher than 90° and circular mean values of V1, LM, RL, and AL regions were smaller than 90° (Fig. 5D). This analysis also showed that there were two clusters according to blood vessel orientation: one was composed of medial regions and the other one was formed by lateral regions and V1. This result agreed well with our THG imaging results where lateral regions and V1 formed a cluster of lower EAL values whereas medial regions formed a cluster of higher EAL values (Fig. 2D).

**Figure 5.**
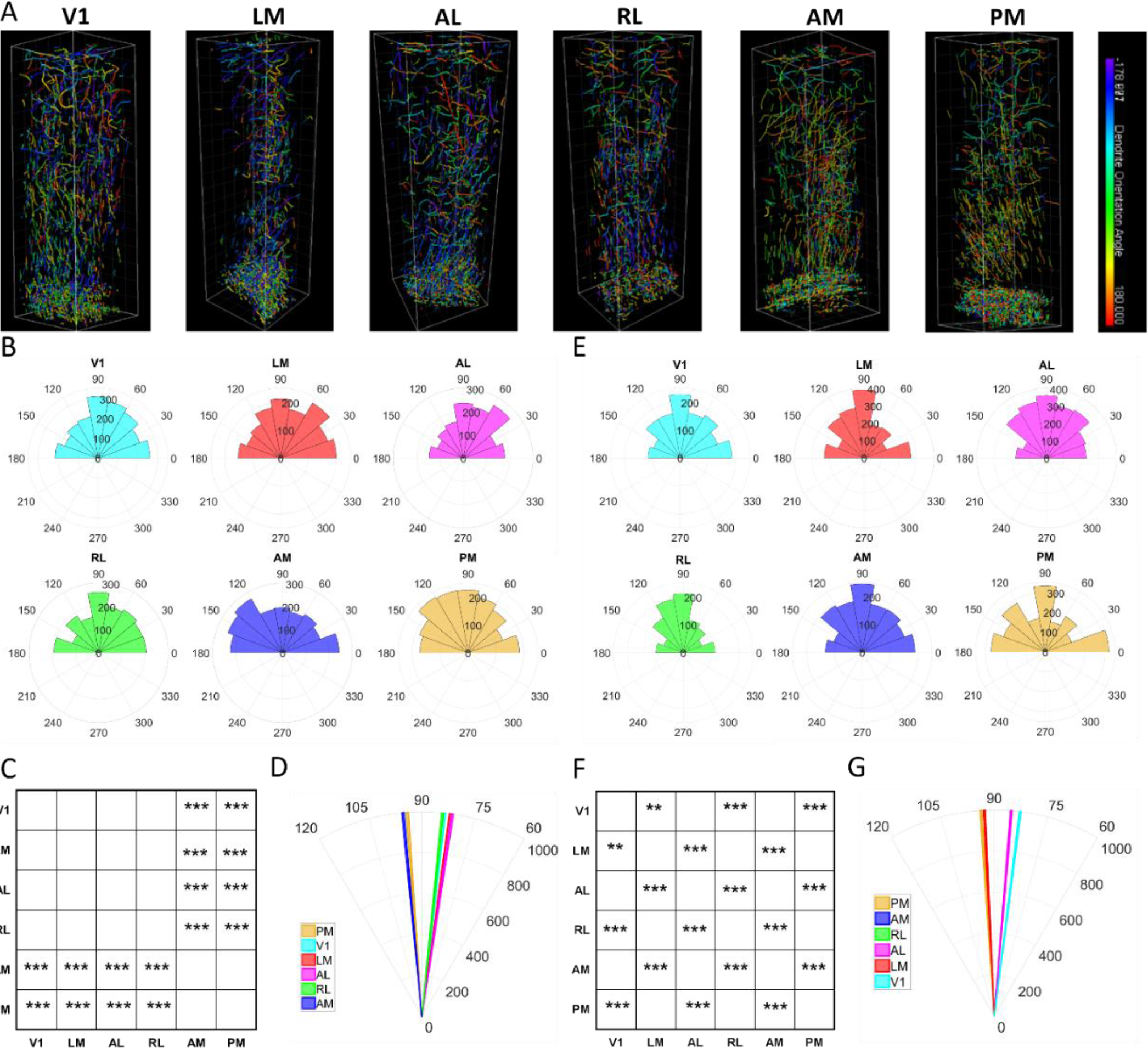
Quantification of orientation of blood vessels and myelin fibers. (A) Three-dimensional depth-resolved THG images for six visual areas highlighting orientation of blood vessels in the cortex and myelin fibers in the white matter. (B) Polar plots of blood vessel orientation in six visual areas. (C) Statistical comparison of circular mean values of blood vessels. (D) Representation of circular mean values of blood vessels in polar plots. (E) Polar plots of myelin fiber orientation in six visual areas. (F) Statistical comparison of circular mean values of myelin fibers. (G) Representation of circular mean values of myelin fibers in polar plots.

We next analyzed the orientation of myelin fibers in the white matter (see Fig 5A) and represented the distribution of their orientation in polar plots for each region (Fig. 5E). Similar to the blood vessel orientation analysis, we applied a circular multi-sample one-factor ANOVA statistical test on the myelin fiber orientation distribution in each visual area (Fig. 5F). This analysis showed that the orientations of myelin fibers in V1, AL, and AM were significantly different from those in LM, RL, and PM. We calculated the circular mean of these orientation distributions and found that circular mean values of LM, RL, and PM were higher than 90° and circular mean values of V1, AL, and AM were smaller than 90° (Fig. 5G). Circular mean values showed that there were two clusters according to myelin fiber orientation: (i) V1, AL, and AM, (ii) LM, RL, and PM. This result also agreed with our THG imaging results where the regions with the same sign in retinotopic maps had similar EAL values: specifically, V1, AL, and AM had higher EAL values, whereas LM, RL, and PM had smaller EAL values.

## Discussion

In this study, we compared EAL values of V1 and higher visual areas by performing label-free THG imaging of blood vessels and myelin fibers in the cortex and in the white matter as well as by performing ablation at four depths in each region. Our first key finding was that most cortical areas had a unique EAL; furthermore, there was a significant difference in EAL values between cortical regions adjacent to each other, and these values exhibited a pattern similar to that of the sign shift that we observed in the retinotopic sign map. These differences can be explained by the following: (i) there may be significant differences in the size, shape, and density of cells (cytoarchitecture of neurons, astrocytes, etc.) in these cortical regions, (ii) the morphology (orientation) of blood vessels and myelin fibers may be different in these regions. Our cytoarchitecture analysis coupled with numerical estimates of EAL values in the cortex showed that there was a strong agreement with our experimental EAL values and their numerical estimates. This result demonstrated that the EAL could be used as a structural metric for a cortical area, which can be obtained via in vivo imaging. Next, we quantified the morphology of blood vessels and myelin fibers with THG imaging for each cortical region, to examine whether there was any relationship between the orientation of blood vessels and myelin fibers in these regions. These data showed that medial regions and lateral regions were significantly different from each other. Interestingly, the orientation of myelin fibers resulted in the same trend as the retinotopic sign map. This result recapitulates the coupling of structure and function of brain regions obtained with diffusion tensor imaging (DTI) of white matter tracts in human and non-human primates^3,42,43^. Overall, the morphology of blood vessels, myelin fibers, and cells showed that morphological differences exist between cortical areas and may lead to significant differences in EAL values of these areas. Structural differences between primary and higher visual areas, including cell size, density^2^, as well as myelin density^42,43^ have been reported in monkeys and humans. To the best of our knowledge, we show for the first time that morphological differences exist in V1 and higher visual areas in mice, in terms of cell size and density in brain sections as well as orientation of blood vessels and myelin fibers obtained in awake conditions. These differences, which likely lead to significant differences between EAL values in these areas, are also reflected in the sign of their retinotopic map.

Why might the retinotopic sign map correlate with structural features of cortex as reflected in the EAL? Adjacent visual areas likely arise from a common border which represents a cardinal axis of visual space (the vertical meridian or the horizontal meridian of the contralateral visual field)^7,44^. During development, thalamocortical fibers may align themselves from this border and follow molecular gradients that confer topography^39,45^. Cortical areas with one sign (V1, AL, AM) may all represent similar thalamocortical (or intracortical) fiber orientations compared to areas with the opposite sign (RL, LM, PM). These fiber orientations would then be reflected in the orientations of myelin and even blood vessels^22,46,47^ as measured with THG imaging that contribute to EALs.

An important finding from our study was that EAL values obtained with ablation experiments were 17 ± 3 % higher than EAL values obtained with THG imaging of blood vessels and myelin fibers. The difference in EAL values obtained with THG experiments and ablation experiments may be attributed to the following: (i) in ablation experiments, we only used the excitation wavelength, whereas in THG imaging we used both the excitation and emission wavelengths to quantify the EAL values, (ii) the interaction between the laser and the brain occurs via optical breakdown as a multi-photon (n>3) phenomenon in ablation, whereas the same interaction occurs as a three-photon phenomenon in THG experiments, (iii) there is a nonuniform distribution of THG signal resources in three-dimensional space that is similar in all imaging modalities used for the quantification of EAL. The main advantage of ablation is that it only relies on the excitation wavelength. However, it requires powerful lasers to create lesions at multiple depths. On the other hand, the main advantage of THG imaging is that it provides realistic EAL values without using any exogenous labeling, but it still has the disadvantage of using both excitation and emission photons to estimate the EAL. To compare our THG imaging results with previous studies based on exogenous labeling of blood vessels, we also performed THG imaging in the somatosensory cortex (S1) of awake mice where the EAL was reported as 315-330 μm^26^. Our THG results revealed that EAL of the S1 was 312.7 ± 2.1 μm in the cortex and 105.4 ± 1.1 μm in the white matter (Supplementary Figure 6). The agreement between our THG imaging results and blood vessel imaging results in the literature^26^ suggests that the EAL of any cortical region in the same animal can be estimated reasonably well by performing label-free THG imaging at 1300 nm excitation wavelength.

## Acknowledgments

This work was supported by US National Institute of Health (NIH) grants EY007023 and EY028219 (MS), 4-P41-EB015871 (PTCS), US National Science Foundation (NSF) grant EF1451125 (MS), a Picower Institute Engineering Collaboration Grant (MS, MY and PTCS), and an equipment grant from the Massachusetts Life Sciences Initiative.

## Methods and Materials

### Surgical Procedures

Experiments were carried out under protocols approved by MIT’s Animal Care and Use Committee and conformed to NIH guidelines. All data in this study were collected from adult (>8 weeks old) mice of either sex. The mouse line was created by crossing Ai93 (TITLa-tTa) with Emx1-IRES-Cre mice lines from The Jackson Laboratory. Mice were initially anesthetized with 4% isoflurane in oxygen and maintained on 1.5-2% isoflurane throughout the surgery. Buprenorphine (1 mg/kg, subcutaneous) and/or meloxicam (1 mg/kg, subcutaneous) was administered preoperatively and every 24 h for 3 days to reduce inflammation. Ophthalmic ointment was used to protect the animal’s eyes during the surgery. Body temperature was maintained at 37.5 °C with a heating pad. The scalp overlying the dorsal skull was sanitized and removed. The periosteum was removed with a scalpel and a craniotomy (5 mm) was made over the primary visual cortex (V1, 4.2 mm posterior, 3.0 mm lateral to Bregma) on either the left or right hemisphere, leaving the dura intact. For calcium imaging, a circular cover glass (5 mm, Warner Instruments) was implanted over the craniotomy as a cranial window, and sealed with dental acrylic (C&B-Metabond, Parkell) mixed with black ink to reduce light transmission. Finally, a custom-designed stainless steel head plate (eMachineShop.com) was affixed to the skull using dental acrylic. Experiments were performed at least 5 days after head plate implantation to allow animals to recover.

### One-photon imaging system for area segmentation in visual areas of awake mice

Widefield one-photon imaging was performed using a custom-made one-photon widefield scope. The cortex was excited by a blue LED light source (Thorlabs) collimated with the objective (NA 0.6). Emitted light was collected with a monochrome CCD camera (1392×1040, spatial resolution 200 pixel per mm with a maximum frame rate of 20 Hz). A large computer display (Fig. 1A) was placed in front of awake head-fixed mice for presenting visual stimuli. To maximize the visual field to be mapped, the plane of display was positioned perpendicular to the plane of the retina at a 30-degree angle relative to the body axis of the animal. To generate a visual retinotopic sign map, animals were exposed to a drifting bar containing flickering black-and-white checkerboard (Fig. 1B). Brain activity was imaged and all six visual areas with their boundaries obtained by the retinotopic map were overlaid on the surface blood vessel image (Fig. 1C-D).

### Custom-made three-photon system

Ultrashort laser pulses (300 fs, 400 kHz, 16 W) at 1045 nm from a pump laser (Spirit, Spectra Physics) were passed through a noncollinear optical parametric amplifier (NOPA, Spectra Physics) to obtain the excitation wavelength of 1300 nm for GCaMP6s and THG imaging. The internal compressor in NOPA could compress the Gaussian pulse width to 22 fs (which was the transform limited pulse width for a 110 nm spectral full-width half-maximum bandwidth laser pulse). Due to the dispersion in the microscope, the Gaussian pulse width on the sample was ~200 fs. In order to shorten the pulse width on the sample, a two-prism-based external compressor was built to prechirp the pulse before sending it to the microscope, reducing the pulse width to 35 fs on the sample. After pulse broadening, a delay line was used to double the repetition rate and increase the frame rate for THG imaging so that a frame rate with 512 × 512 pixels for a 350 × 350 μm^2^ field of view could be increased up to 2 Hz. Power control was performed with the combination of a half-wave plate (AHWP05M-1600, Thorlabs) and a low-GDD ultrafast beamsplitter (UFBS2080, Thorlabs) with 100:1 extinction ratio. The laser beams were scanned by a pair of galvanometric mirrors (6215H, Cambridge Technologies) to image the laser spot on the back aperture of the objective using a pair of custom-designed scan and tube lenses. The emitted signal from the mouse brain was collected by a pair of collection lenses for three PMTs. GCaMP6s fluorescence signals were detected using GaAsP photomultiplier tubes (H7422A-40, Hamamatsu, Japan); THG signal was detected using a bialkali (BA) photomultiplier tube (R7600U-200). A commercially available objective with high transmission at a longer wavelength (25x, 1.05-NA, XLPN25XWMP2, Olympus) was used. Image acquisition was carried out using ScanImage (Vidrio). Imaged cells, blood vessels, and myelin fibers were located at a depth of 0-1200 μm or more below the pial surface. Laser power ranged from 0.5-50 mW at the sample depending on depth and fluorescence expression levels. Awake mice were placed on a two-axis motorized stage (MMBP, Scientifica) and the objective lens was placed on a single axis motorized stage (MMBP, Scientifica) to move in the axial direction. Mice were fixed on the stage with a sample holder, and a head mount was placed on top of the head to minimize motion artifacts during imaging.

### Laser ablation experiments

To determine the EAL of the visual cortex and other visual areas, laser ablation at 1300 nm at 1 kHz repetition rate was performed. The laser beams were raster scanned at a single depth below the pia surface for the duration of one frame, namely 20 s (0.05 fps), using a pair of galvanometric mirrors. For the targeted ablation field of view of 50 μm and 512 × 512 pixel rate, minimum number of overlapping pulses resulted in 10 s of ablation duration with a 1.3 μm 1/e^2^ one-photon lateral resolution.

Since EAL is defined as an effective attenuation length for each region, we can model light attenuation in the brain with EAL and Beer-Lambert’s law as follows:

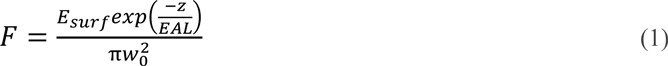

where E_surf_ is the laser energy at the surface, z is the ablation depth, EAL is the effective attenuation length of cortex, w_0_ is the one-photon 1/e^2^ radius at the focal plane which can be calculated from the three-photon point-spread function^19^, and F is the fluence at the focal plane. To calculate the threshold fluence and EAL, Eq. 1 can be reorganized as follows:

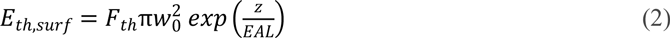

By taking the natural logarithm of both sides in Eq. 2, a linear regression can be applied to calculate F_th_ and EAL:

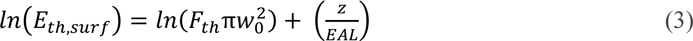

Threshold energy (E_th,surf_) for each depth is calculated by fitting percent of damage with respect to applied energy on the surface. This fitting function is represented as follows:

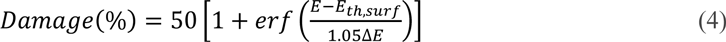

After determining threshold energy for each depth, a linear regression was applied to ablating four different depths in V1 and each of the other visual areas.

### Calculation of combined EAL for the cortex and the white matter

After calculating EALs for the cortex and the white matter, we can define a single EAL (EAL_comb_) which combines these two EALs as follows:

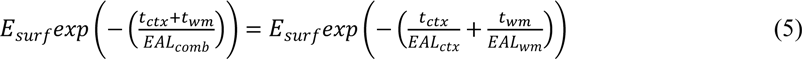

where t_ctx_ and t_wm_ are the thicknesses of the cortex and the white matter, respectively. Similarly, EAL_ctx_ and EAL_wm_ are the EALs of the cortex and the white matter, respectively. To calculate the EAL_comb_, Eq. 5 can be reorganized as follows:

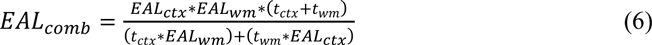

### Quantification of cell size and density

Low magnification images of brain slices were taken with a 10x magnification objective (Olympus). We performed tiling in lateral directions, and depth-resolved imaging with 10 μm increment in the axial direction, followed by maximum intensity projection to register the final image to the Allen Brain Atlas using a custom algorithm^40^. Then, we located each visual area and acquired high magnification images with a 20x magnification objective (Olympus) with 1 μm increment in the axial direction. Finally, we performed maximum intensity projection and applied tha “Analyze Particles” module in Image J to quantify the cell size and cell density for each visual area.

### Relationship between scattering length and cytoarchitecture of visual areas

From Mie theory, the scattering length (*l*_*s*_) of biological tissues can be calculated as follows^21^:

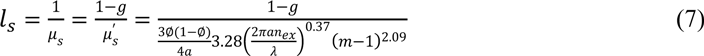

where *μ*_*s*_ is scattering coefficient, *μ*′_*s*_ is reduced scattering coefficient, *g* is average cosine of the scattering angle, ∅ is the volume fraction or cell density, *a* is the radius of the cell, *λ* is the wavelength of the light, *m* = (*n*_*in*_/*n*_*ex*_) is the refractive index of the intracellular and extracellular fluid, respectively. We can calculate scattering length of each visual area by taking *n*_*ex*_ = 1.33 and m= 1.04 as determined previously^21^. Then, we can combine the scattering lengths of each visual area with the absorption length of water (*l*_*a*_) at 1300 nm^41^ to calculate EAL values as follows:

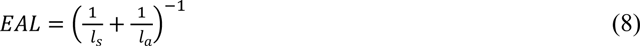

### Quantification of orientation of blood vessels and myelin fibers

To quantify the orientation of blood vessels in the cortex and the orientation of myelin fibers in the white matter, we performed depth-resolved THG imaging with 1 μm increment in the axial direction. With this fine incremental imaging, we could calculate the orientation of small blood vessels and myelin fibers via Imaris software. Imaris software provided an orientation map spanning from −180° to +180°; we converted this map to a range of 0 to +180°.

### Image analysis for ablation experiments

An image analysis algorithm was developed to calculate percent of damage in the GCaMP6s images. First, a median filter with a size of 2 × 2 was applied to smooth the images. Otsu’s thresholding method^48^ was applied to remove background noise and to convert the images into binary ones. Then, the percent of damage was calculated by taking the ratio of difference of total number of pixels below the threshold before and after in ablation images to the total number of pixels corresponding to the targeted ablation area.

### Statistics

EAL values in cortex and white matter of visual areas were compared using one-way ANOVA with multiple comparisons (Fig. 3). One-factor ANOVA (or Watson-Williams test) in MATLAB circular statistics toolbox was applied to the data presented in Figure 5. Specifically, we compared two visual areas as to whether they have similar or different circular mean values (Figs. 5C and 5F), and one visual area with multiple visual areas such as V1 to AM and PM (Fig. 5C) or V1 to LM, RL, and PM (Fig. 5F).

### Histology

Animals were deeply anesthetized with 4% isoflurane and perfused transcardially with 0.1 M phosphate-buffered saline (PBS) followed by chilled 4% paraformaldehyde in 0.1 MPBS. The brains were then postfixed in 4% paraformaldehyde in 0.1 M PBS (<4 °C) overnight. The fixed brains were sectioned into 100 μm slices with a vibratome and then mounted on a glass slide with Vectashield Hardset Antifade Mounting Medium with DAPI (Vector Labs). The slides were imaged using a confocal microscope (Leica TCS SP8).

## Supplementary Figures

**Supplementary Figure 1.**
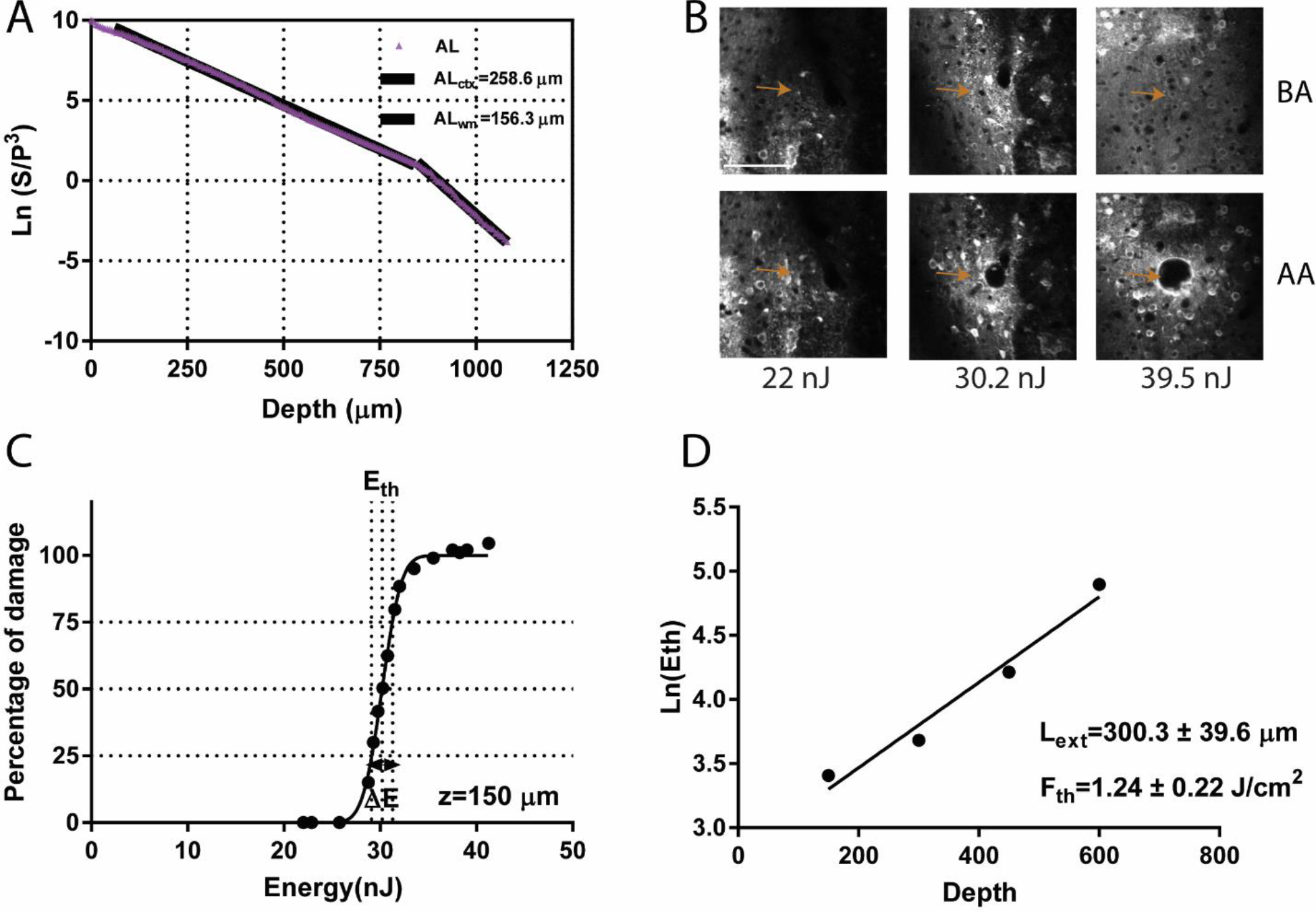
Characterization of EAL values via THG imaging and ablation in AL. **(A)** THG imaging in AL results in 258.6 ± 10.4 μm effective attenuation length for the cortex and 156.3 ± 9.5 μm effective attenuation length for the white matter. **(B)** Representative images before ablation (BA) and after ablation (AA) for three pulse energies (22, 30.2 and 39.5 nJ) at 150 μm depth. Arrows show the location of target region for the ablation before and after applying laser pulses for each pulse energy. (C) Determining extinction length via tissue ablation. Percent of damage ranges from 0 to 100% with respect to laser energy on the tissue surface. Threshold energy (*E*_th_) is the energy which results in 50% damage. For ablation at 150 μm depth, *E*_th_ is 30.2 nJ. **(D)** Semi-logarithmic plot of threshold energies for 4 different depths results in attenuation length of 300.3 ± 39.6 μm and threshold fluence of 1.24 ± 0.22 J/cm^2^.

**Supplementary Figure 2.**
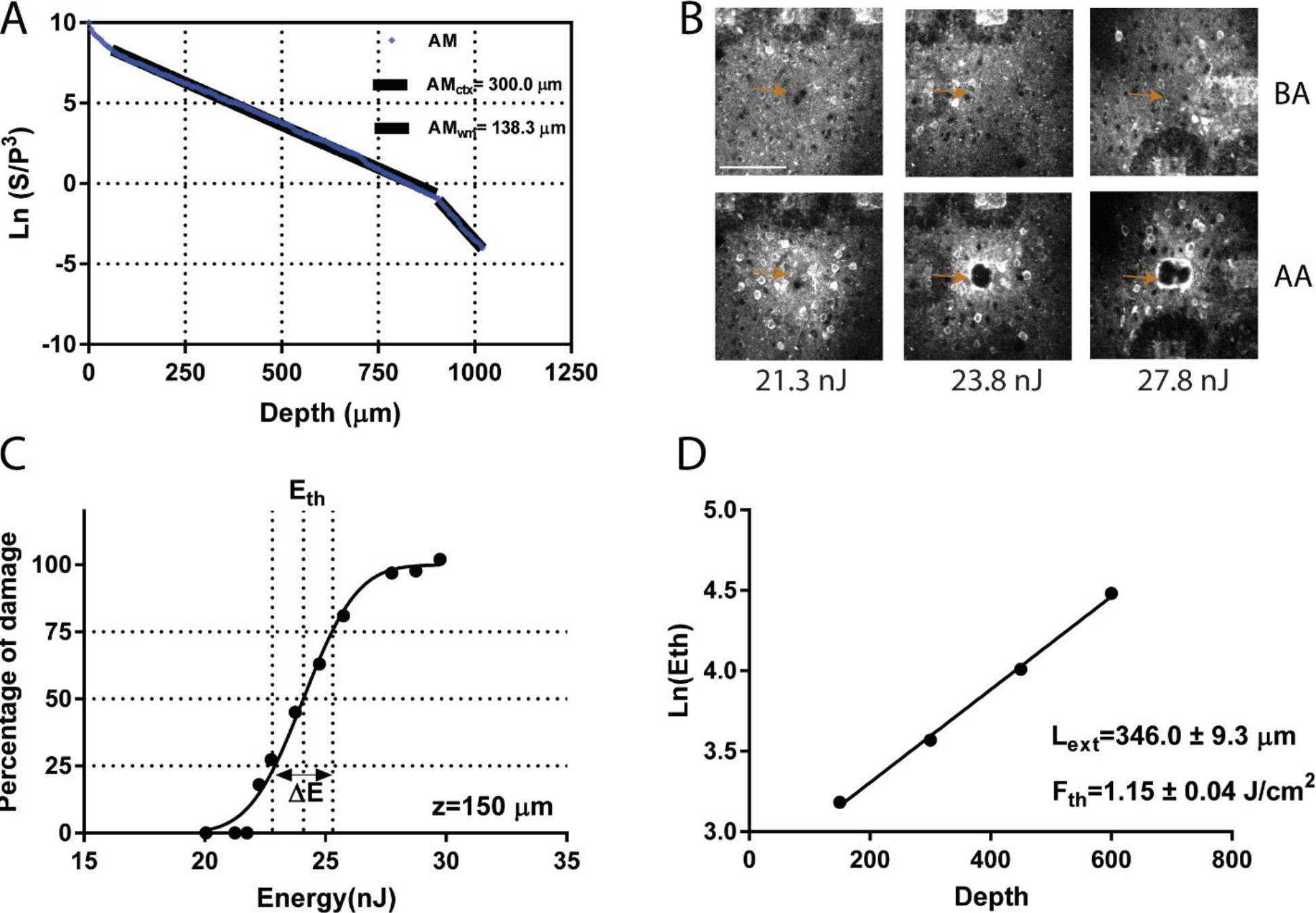
Characterization of EAL values via THG imaging and ablation in AM. **(A)**THG imaging in AL results in 300.0 ± 11.9 μm effective attenuation length for the cortex and 138.3 ± 4.5 μm effective attenuation length for the white matter. **(B)** Representative images before ablation (BA) and after ablation (AA) for three pulse energies (21.3, 23.8 and 27.8 nJ) at 150 μm depth. Arrows show the location of target region for the ablation before and after applying laser pulses for each pulse energy. (C) Determining extinction length via tissue ablation. Percent of damage ranges from 0 to 100% with respect to laser energy on the tissue surface. Threshold energy (*E*_th_) is the energy which results in 50% damage. For ablation at 150 μm depth, *E*_th_ is 24.1 nJ. **(D)** Semi-logarithmic plot of threshold energies for 4 different depths results in attenuation length of 346.0 ± 9.3 μm and threshold fluence of 1.15 ± 0.04 J/cm^2^.

**Supplementary Figure 3.**
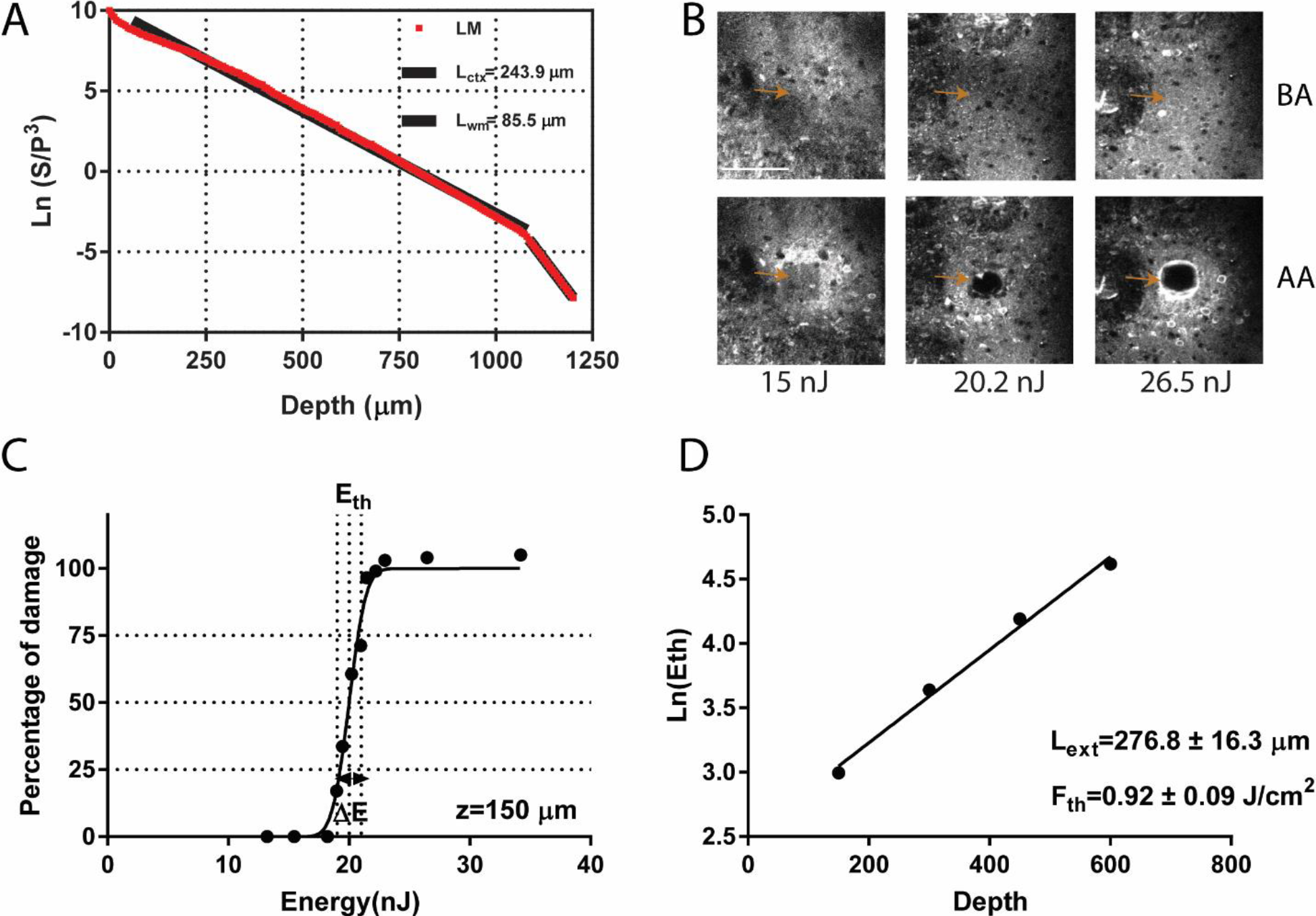
Characterization of EAL values via THG imaging and ablation in LM. **(A)** THG imaging in AL results in 243.9 ± 11.2 μm effective attenuation length for the cortex and 85.5 ± 3.5 μm effective attenuation length for the white matter. **(B)** Representative images before ablation (BA) and after ablation (AA) for three pulse energies (15, 20.2 and 26.5 nJ) at 150 μm depth. Arrows show the location of target region for the ablation before and after applying laser pulses for each pulse energy. (C) Determining extinction length via tissue ablation. Percent of damage ranges from 0 to 100% with respect to laser energy on the tissue surface. Threshold energy (*E*_th_) is the energy which results in 50% damage. For ablation at 150 μm depth, *E*_th_ is 20 nJ. **(D)** Semi-logarithmic plot of threshold energies for 4 different depths results in attenuation length of 276.8 ± 16.3 μm and threshold fluence of 0.92 ± 0.09 J/cm^2^.

**Supplementary Figure 4.**
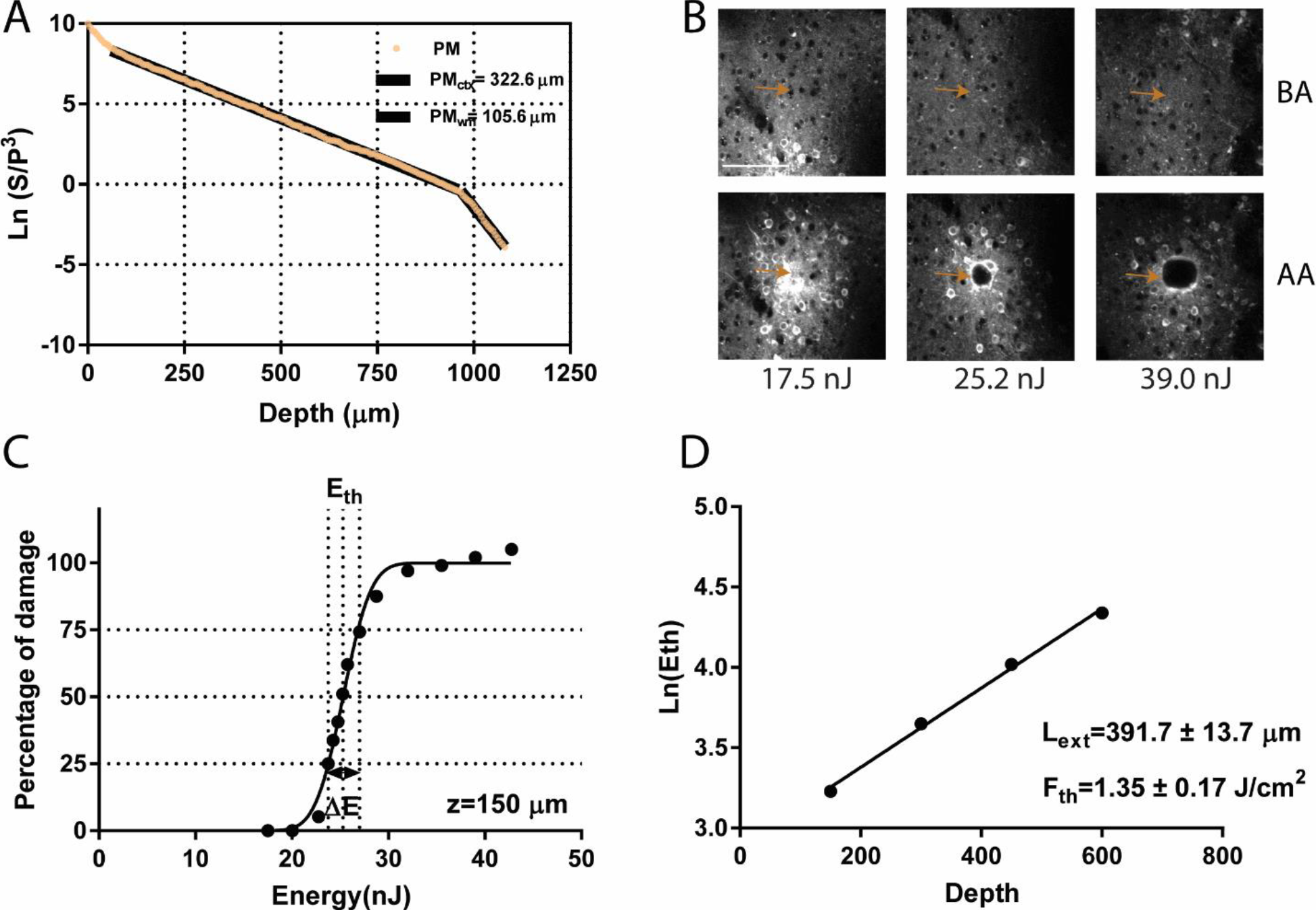
Characterization of EAL values via THG imaging and ablation in PM. **(A)** THG imaging in AL results in 322.6 ± 14.5 μm effective attenuation length for the cortex and 105.6 ± 4.3 μm effective attenuation length for the white matter. **(B)** Representative images before ablation (BA) and after ablation (AA) for three pulse energies (17.5, 25.2 and 39.0 nJ) at 150 μm depth. Arrows show the location of target region for the ablation before and after applying laser pulses for each pulse energy. (C) Determining extinction length via tissue ablation. Percent of damage ranges from 0 to 100% with respect to laser energy on the tissue surface. Threshold energy (*E*_th_) is the energy which results in 50% damage. For ablation at 150 μm depth, *E*_th_ is 25.3 nJ. **(D)** Semi-logarithmic plot of threshold energies for 4 different depths results in attenuation length of 391.7 ± 13.7 μm and threshold fluence of 1.35 ± 0.17 J/cm^2^.

**Supplementary Figure 5.**
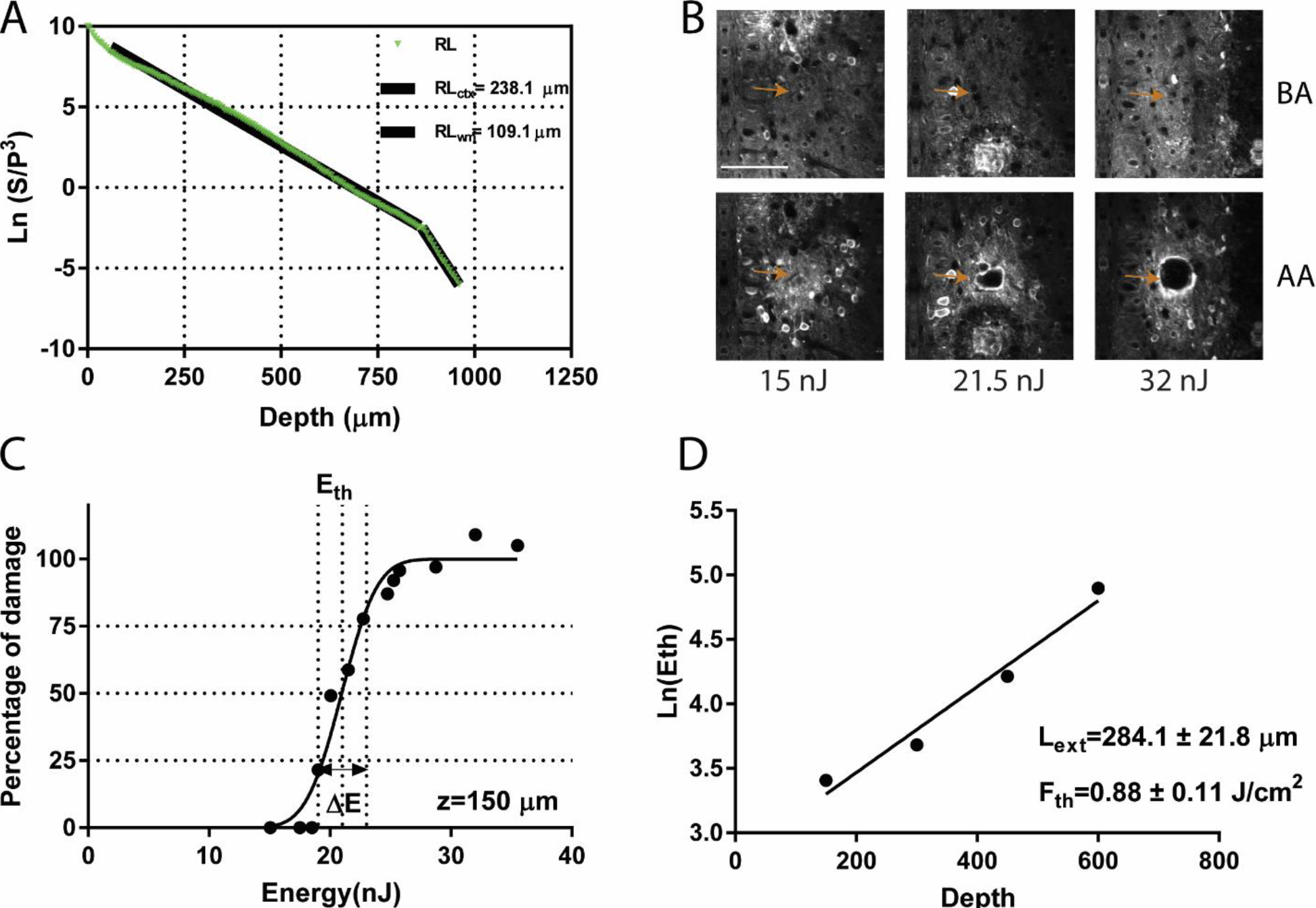
Characterization of EAL values via THG imaging and ablation in RL. **(A)** THG imaging in AL results in 238.1 ± 14.0 μm effective attenuation length for the cortex and 109.1 ± 2.9 μm effective attenuation length for the white matter. **(B)** Representative images before ablation (BA) and after ablation (AA) for three pulse energies (15, 21.5 and 32.0 nJ) at 150 μm depth. Arrows show the location of target region for the ablation before and after applying laser pulses for each pulse energy. (C) Determining extinction length via tissue ablation. Percent of damage ranges from 0 to 100% with respect to laser energy on the tissue surface. Threshold energy (*E*_th_) is the energy which results in 50% damage. For ablation at 150 μm depth, *E*_th_ is 21.0 nJ. **(D)** Semi-logarithmic plot of threshold energies for 4 different depths results in attenuation length of 284.1 ± 21.8 μm and threshold fluence of 0.88 ± 0.11 J/cm^2^.

**Supplementary Figure 6.**
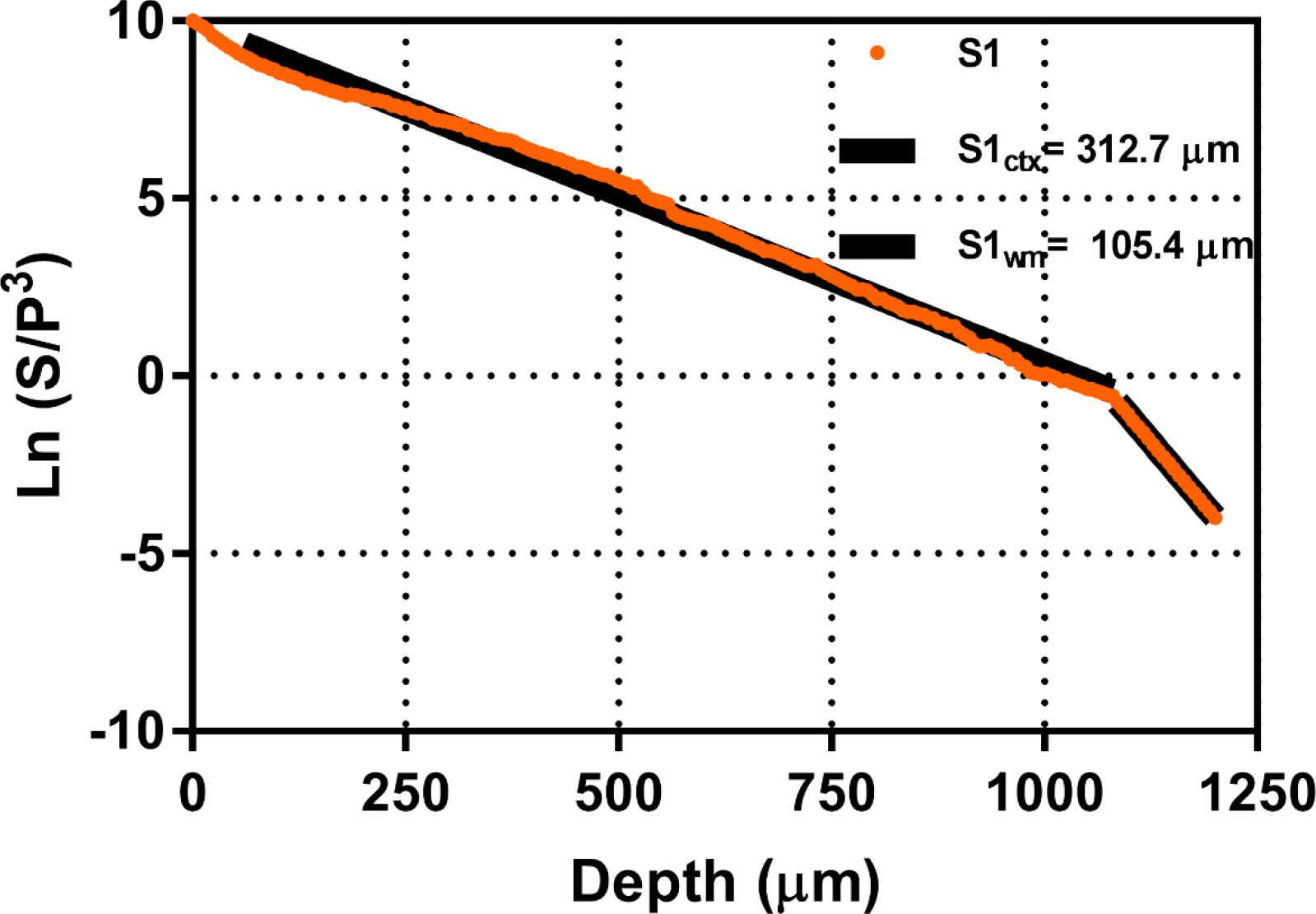
Characterization of EAL values via THG imaging in S1. THG imaging in S1 results in 312.7 ± 2.1 μm effective attenuation length for the cortex and 105.4 ± 2.1 μm effective attenuation length for the white matter. Raw data is in orange, linear fits are in black.

